# The non-genomic vitamin D pathway links β-amyloid to autophagic apoptosis in Alzheimer’s disease

**DOI:** 10.1101/2021.05.06.443028

**Authors:** Rai-Hua Lai, Yueh-Ying Hsu, Feng-Shiun Shie, Mei-Hsin Chen, Jyh-Lyh Juang

## Abstract

Vitamin D is an important hormonal molecule, which exerts genomic and non-genomic actions in maintaining brain development and adult brain health. Many epidemiological studies have associated vitamin D deficiency with Alzheimer’s disease (AD). Nevertheless, the underlying signaling pathway through which this occurs remains to be characterized. We were intrigued to find that although vitamin D levels are significantly low in AD patients, their hippocampal vitamin D receptor (VDR) levels are inversely increased in the cytosol of the brain cells, and colocalized with Aβ42 plaques, gliosis and autophagosomes, suggesting that a non-genomic form of VDR is implicated in AD. Mechanistically, Aβ42 induces the conversion of nuclear heterodimer of VDR/RXR heterodimer into a cytoplasmic VDR/p53 heterodimer. The cytosolic VDR/p53 complex mediates the Aβ42–induced autophagic apoptosis. Reduction of p53 activity in AD mice reverses the VDR/RXR formation and rescues AD brain pathologies and cognitive impairment. In line with the impaired genomic VDR pathway, the transgenic AD mice fed a vitamin D sufficient diet exhibit lower plasma vitamin D levels since early disease phases, raising the possibility that vitamin D deficiency may actually be an early manifestation of AD. Despite the deficiency of vitamin D in AD mice, vitamin D supplementation not only has no benefit but lead to exacerbated Aβ42 depositions and cognitive impairment. Together, these data indicate that the impaired genomic vitamin D pathway links Aβ42 to induce autophagic apoptosis, and suggest that VDR/p53 pathway could be targeted for the treatment of AD.

**Significance Statement:** Vitamin D exerts a genomic action for neuroprotection through VDR/RXR transcriptional complex. Thus, insufficient vitamin D has been linked to AD, but the signaling pathway involved remains unclear. Surprisingly, we find that the genomic action of VDR/RXR to be compromised and converted into a non-genomic VDR/p53 complex in promoting AD neurodegeneration. The cytosolic VDR/p53 complex contribute to autophagy-induced neuronal apoptosis. The VDR/RXR pathway can be a new therapeutic target for AD because targeting VDR/p53 ameliorates AD. Importantly, we provide evidence that vitamin D deficiency might be an early AD manifestation, and vitamin D supplementation exacerbates AD. This work uncovers a non-genomic VDR action in promoting AD and suggests a potential aggravating effect of vitamin D supplementation on AD.

## Introduction

Dementia is increasing in prevalence as the world’s populations age (1). Alzheimer’s disease (AD) accounts for 60 to 80 percent of all cases of dementia(2, 3), but no therapy has been found to prevent or decelerate progression of the disease. Thus, it is important to identify early modifiable risk factors to prevent or treat this debilitating disease. Many epidemiological studies have suggested that vitamin D deficiency could be a potential risk factor for AD and other age-related chronic diseases (4). Vitamin D is actually not a vitamin; but a steroid hormone (5). In addition to its well-known contribution to mineral and skeletal homeostasis, it also plays neurotrophic or neuroprotective roles in brains (6). Like other steroid hormone, vitamin D exerts both genomic and nongenomic effects on human brain. The genomic action of vitamin D is initiated via binding to vitamin D receptor (VDR). In specific, the vitamin D metabolite 1α,25-dihydroxyvitamin D3 (or calcitriol) binds to VDR. The ligand-bound VDR prefers dimerization with retinoid X receptor (RXR) for transcription regulation of a set of genes containing vitamin D-response elements (VDREs) in their promoter/regulatory regions. For instance, CYP24A1, a key vitamin D catabolism enzyme, is a typical VDRE-containing target gene that is positively regulated by VDR (7, 8). There is abundant evidence that vitamin D exerts genomic actions in the brain. In supporting this concept, VDR is found to be widely expressed in major brain cell types in mediating brain development and function (9).

Besides the classical genomic activity, vitamin D also exerts non-genomic actions on the brain but it is less well characterized (10, 11). In contrast to genomic action, the non-genomic action of VDR is a rapid plasma membrane response to cellular stimuli. However, the non-genomic action of VDR does not seem to require the VDR-RXR interaction. Of special note is the finding that VDR is also involved in xenobiotic metabolism in detoxification processes (12, 13), which is independent of vitamin D binding (14). Since soluble Aβ protofibrils are assumed to mediate the neurodegeneration in AD (15), it is of great importance to investigate whether VDR signaling is involved in this Aβ-induced toxic effect.

Previously, assuming that vitamin D was protective, we were surprised that deficiencies in that vitamin does not seem to produce the same AD symptoms as deficiencies in vitamin B12 or folate in humans (16-18). Thus, we became interested in investigating whether VDR levels to be decreased in the AD brains due to the deficiency of it canonical ligand. We then performed a series of mechanistic investigations studying the VDR expression and distribution in the AD brains and the role of VDR/p53 complex formation in the AD pathogenesis. To determine whether VDR/p53 complex was involved in the promotion of AD pathogenesis, we applied a chemical inhibitor of p53 on AD mice. To answer the question of whether or not vitamin D deficiency to be an early manifestation of AD, we studied the effect of a vitamin D sufficient diet on serum levels of vitamin D in APP/PS1 AD and wild-type (WT) mice. Finally, we studied the effect of vitamin D supplementation on the disease progression in AD mice to interrogate the proposition that vitamin D supplementation may exert a role in protecting AD. The results support the critical role of the non-genomic action of cytosolic VDR/p53 complex in inducing atuophagic apoptosis and may represent a novel therapeutic target for AD.

## Results

### Increased brain VDR levels in human and mouse with AD

Since VDR functions as a ligand-activated transcription factor upon vitamin D binding, we anticipated to observe inhibition of VDR in response to the deficiency of its canonical ligand in the AD brains. We studied hippocampal tissues obtained postmortem from AD patients to find out whether the same phenomenon occurred in humans with AD. Performing Western blot and immunohistochemistry analyses, we found marked increases in VDR proteins in these tissues (Fig. 1*A*, 1*B* and SI Appendix, Fig. S1). Similar phenomenon was also observed in APP/PS1 mouse brains. Of special note is the older the mouse, the more evident the increase in VDR (Fig. 1*C* and SI Appendix, Fig. S2*A*). In addition, this increase in VDR proteins was found to widely distribute among various cell types, including neurons, astrocytes and microglia (Fig 1*D* and SI Appendix, Fig. S2*B*). Of special note, the increased VDR proteins were predominantly localized to the cytosolic fraction (Fig. 1*D*).

**Figure 1.**
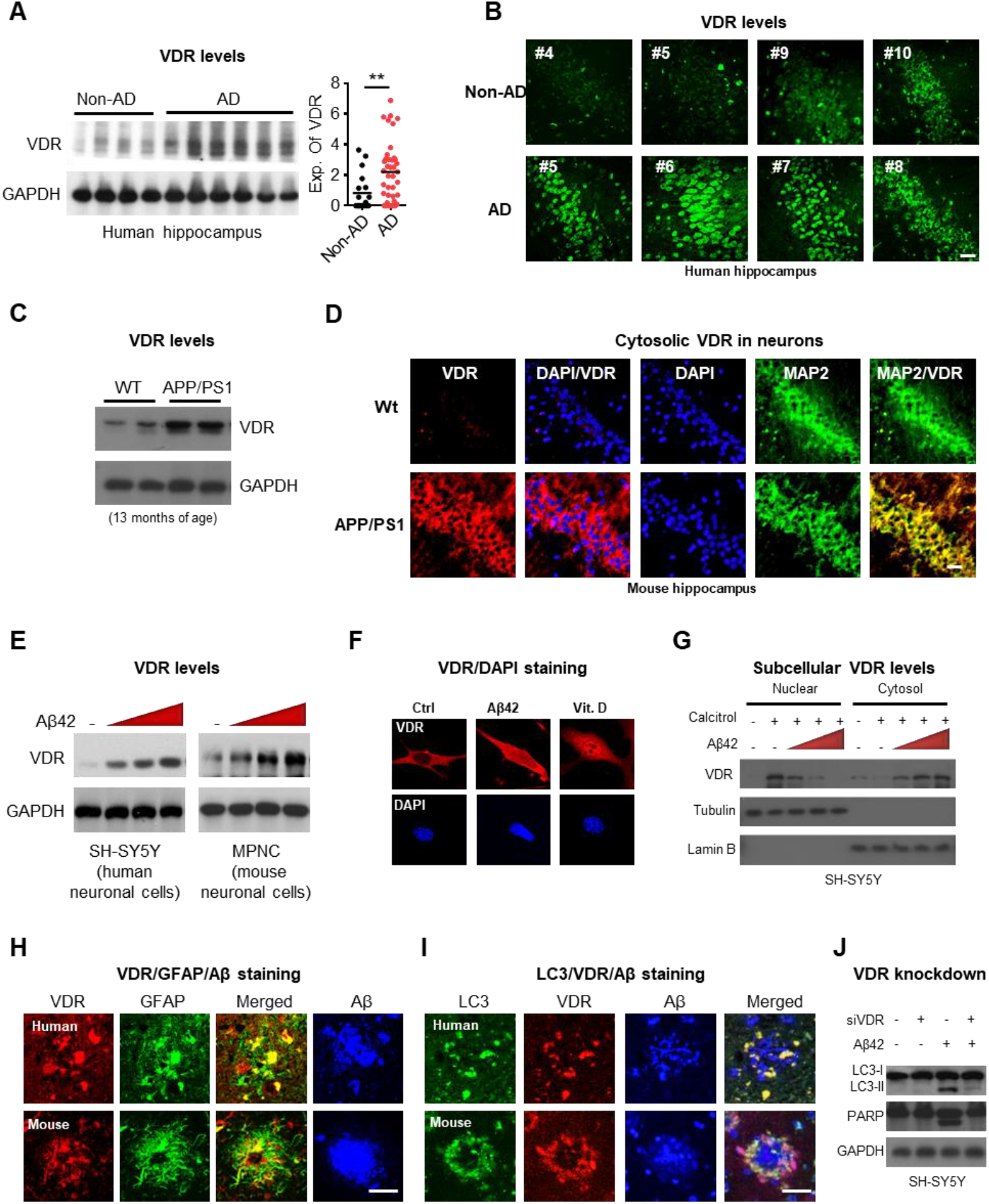
A non-genomic activation of VDR proteins in human and mouse brains with AD. *(A)* Western blot analysis of VDR levels in the hippocampal tissue lysates of AD patients (AD) and age-matched non-AD controls (Non-AD). Densitometrical quantification of VDR as ratios normalized with GAPDH (based on Supplemental Fig. S1; AD n=40; non-AD n=18). **P < 0.01 by unpaired t-test (right panel). *(B)* Representative immunohistochemistry analysis of VDR in sections of CA regions in AD and non-AD controls. *(C)* Western blot analysis of VDR levels in hippocampus of APP/PS1 and wild type (WT) mice at 13 months of age. *(D)* Representative images of immunohistochemistry analysis for MAP2 (neuronal marker) and VDR in sections of CA regions of hippocampus in APP/PS1 mice. *(E)* Dose-dependent response of VDR protein expressions to Aβ42 treatment in neuronal cells. SH-SY5Y and MPNC cells were treated with 0, 2, 4 or 6 μM Aβ42 for 6 h prior to western blot analysis of VDR protein levels. *(F)* Representative immunofluorescent staining shownβ-amyloid treatment increases predominantly cytosolic VDR protein level. *(G)* VDR levels in cytoplasmic and nuclear fractions determined by Western blot analysis. SH-SY5Y cells were treated with 100 nM calcitriol and 0, 2, 4 or 6 μM Aβ42 for 6 h prior to assays. *(H)* Representative immunofluorescent micrographs of VDR/GFAP/Aβ triple staining in the hippocampal tissue sections of human and mouse AD brains. Scale bars, 20 μm. *(I)* Representative immunofluorescent micrographs of LC3/VDR/Aβ triple staining in the hippocampal tissue sections of human and mouse AD brains. Scale bars, 20 μm. *(J)* Western blot analysis of LC3 and PARP cleavage level in SH-SY5Y cells with or without RNAi knockdown of VDR. SH-SY5Y cells were treated with or without RNAi knockdown of VDR, and then treated with Aβ (6 μM) for 6 h before harvesting for western blot.

Since the observed brain VDR protein levels appeared to be inversely upregulated under the deficiency of vitamin D in AD patients, it was of interest to investigate whether this was a cellular response to xenobiotic stimulus of Aβ because VDR has been known to implicate in endobiotic/xenobiotic-activated metabolism in a vitamin D independent manner (12). We found that by exposing cultured neuronal cells to Aβ42 we could increase VDR protein levels in a dose-dependent manner in the absence of its canonical ligand vitamin D (Fig. 1*E*). To determine whether the Aβ-induced VDR also localized to cytosol as that observed in AD brains, we conducted immunofluorescence staining and also prepared fractionated cytoplasmic and nuclear lysates for Western blot assay. Both assays clearly showed that the increased VDR proteins were largely retained in the cytosol upon Aβ42 stimulus (Fig. 1*F* and 1*G*), supporting our observation of VDR cytosolic localization in AD brains. These results of increased VDR in cytosol may suggest a non-genomic activation of VDR in AD.

### Increased VDR proteins are localized to autophagosome, Aβ plaques and reactive gliosis

To find out how VDR might be involved in pathogenesis of AD at the cellular level, we first investigated whether the activated VDR might be associated with specific pathological features of AD. We found the autophagy marker LC3, reactive gliosis, and Aβ plaques to be most pronounced in the hippocampal lesions of AD subjects where VDR signals were amplified (Fig. 1*H*, 1*I* and SI Appendix, Fig. S3). Since autophagy is a dysregulated process reportedly crucial in Aβ pathology, we explored how VDR might be involved in pathogenesis of AD at the cellular level. We knocked down VDR in neuronal cells and found it abrogated Aβ-induced autophagy and apoptosis (Fig. 1*J*), indicating that VDR upregulation was prerequisite to the induction of neuronal autophagy. Collectively, our studies of human brain sections, transgenic mice, and cells highlight a potential detrimental role that VDR plays in transducing Aβ neurotoxic signaling in AD.

### Aβ induces disruption of VDR/RXR and formation of VDR/p53

We wanted to identify the potential molecular mechanisms underlying the non-genomic activation and regulation of the VDR pathway in AD. In the genomic signaling pathway, vitamin D_3_ stimulates VDR to form a complex with retinoid X receptor (RXR), which is then imported into the nucleus for transcription of many target genes (19). However, our studies of hippocampal neurons of AD mice as well as in Aβ-treated neuronal cells showed that the enhanced signal of VDR to be largely retained in the cytoplasmic compartment and not translocated into the nucleus. In accordance with these observations, we also found that the VDR/RXR heterodimer did not form in Aβ-treated SH-SY5Y cells (Fig. 2*A*), suggesting possible Aβ disablement of the VDR-RXR pathway. Supporting this idea, the vitamin D_3_-induced VDR/RXR interaction was also found to be dose-dependently abrogated by Aβ42 (Fig. 2*B*). Moreover, the SH-SY5Y cells treated with Aβ42 showed no induction of Cyp24a1, a canonical target gene of VDR, compared with a vitamin D_3_ positive control (Fig.2*C*). Thus, we directly assessed VDR transcriptional activity by reporter assay and found that the Aβ-induced VDR protein did not induce target gene Cyp24a1 expression in neuronal cells, while vitamin D_3_ did (Fig.2*D*). Considered together, these results suggest that Aβ impairs the VDR-RXR pathway in AD.

**Figure 2.**
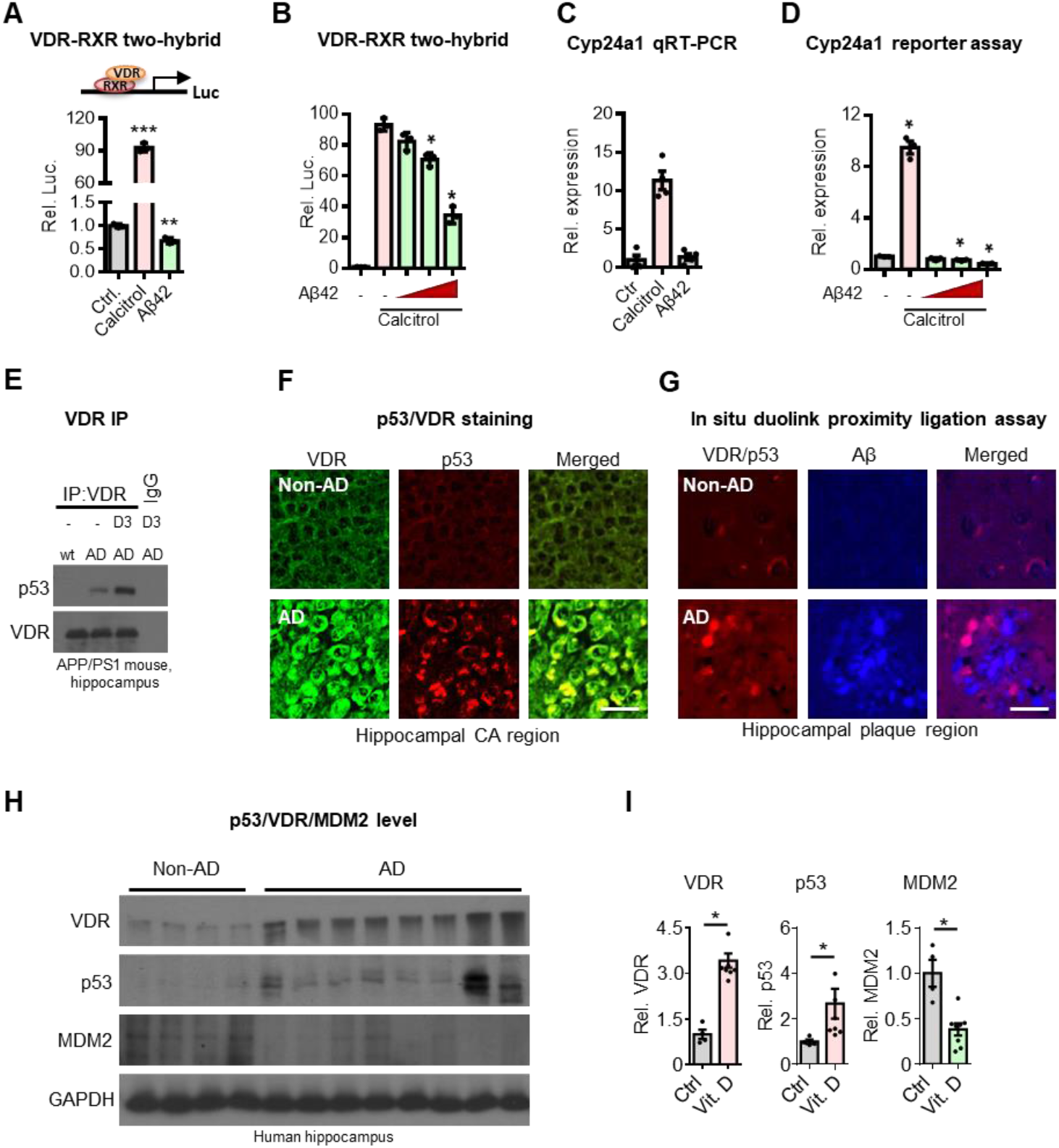
Aβ42 disrupts genomic VDR/RXR complex but induces the formation of VDR/p53 non-genomic complex. *(A)* Mammalian two-hybrid assays for studies of interaction of VDR with RXR in neuronal cells exposed to Aβ42. SH-SY5Y cells were treated with 100 nM calcitriol or Aβ42 for 6 h before harvesting for mammalian two-hybrid luciferase assays (n=3). Values are represented as the mean ± SEM and *P < 0.05, **P < 0.01 or ***P<0.001 by unpaired t-test. *(B)* Aβ42 decreases VDR-RXR interaction in a dose-dependent manner. SH-SY5Y cells were treated with 100 nM calcitriol and Aβ42 (2, 4 or 6 μM) for 6 h before harvesting for mammalian two-hybrid luciferase assays (n=3). *(C)* Decreased Cyp24a1 (a known VDR target gene) gene expression in neuronal cells exposed to Aβ42. SH-SY5Y neuronal cells were treated with 100 nM calcitriol and/or 4 μM Aβ42 for 6 h prior to qPCR analysis of the Cyp24a1 expressions (n=3). *(D)* Cyp24a1 promoter reporter assay in neuronal cells exposed to Aβ42. SH-SY5Y cells were treated with 100 nM calcitriol or Aβ42 (2, 4 or 6 μM) for 6 h before harvesting for luciferase assays (n=3). *(E)* Western blot analysis of co-immunoprecipitated VDR and p53 in the hippocampal tissue lysates of APP/PS1 mouse supplemented with or without of cholecalciferol. *(F)* Representative immunofluorescent micrographs showing p53 and VDR co-localized in hippocampal tissues of human AD brains. Hippocampal CA regions of AD patients were labelled with antibodies against VDR and p53. Scale bars, 20 μm.*(G)* Duolink® proximity ligation assay for protein interaction between VDR and p53 in human hippocampal tissues. Scale bars, 20 μm. (H) Concomitant upregulation of VDR and p53 proteins in the hippocampal tissues of AD. Western blot analysis of hippocampal tissue lysates from AD patients and normal controls with antibodies against p53, VDR, MDM2 and GAPDH (left panel). Densitometrical quantification of the indicated proteins as ratios normalized with GAPDH. *P < 0.05 by unpaired t-test (right panel).

VDR usually forms heterodimers with other transcription factors but we found no interaction between VDR and RXR, so we began to wonder whether VDR switched its interacting partnership with RXR to another transcription factor. P53 has been shown to interact with and modulate VDR activity in tumor cell apoptosis (20), and most importantly, p53 protein has been found to be upregulated in AD brains (21). Therefore, we wanted to test whether VDR switched its partnership with RXR to p53. Indeed, we found this VDR/p53 interaction in AD mice but not WT mice (Fig. 2*E*). Supporting this, double-labelling immunocytochemistry showed that both VDR and p53 were colocalized in cytoplasm (Fig. 2*F*). To validate this result in human brains, we performed a proximity ligation assay (PLA) to study the association between VDR and p53 in the postmortem brains of AD patients. As we had found in mice, VDR/p53 complex was abundant in the plaque regions in the brain sections obtained from AD patients (Fig. 2*G* and SI Appendix, Fig. S4). Moreover, Western blot analysis also suggested that both VDR and p53 protein levels were increased while the MDM2, an enzyme that targets p53 to degradation by ubiquitination, levels were concomitantly decreased in human brains (Fig. 2*H* and 2*I*).

To determine whether the formation of VDR/p53 complex was induced by the direct cellular impact of Aβ42, we performed co-immunoprecipitation assays and found that Aβ triggered VDR/p53 interaction in SH-SY5Y cells and that this interaction was further enhanced with the addition of vitamin D_3_ (SI Appendix, Fig. S5*A*). Consistently with the observations in AD brain tissues, the Western blot results also showed that both the VDR and p53 proteins were also predominantly localized within the cytosolic compartment in SH-SY5Y cells exposed to Aβ42 (SI Appendix, Fig. S5*B*). Given these data together, we reasoned that the resulting VDR/p53 protein complex might exert non-genomic actions on AD brains because the p53 and VDR proteins were found to be largely localized to the cytoplasm in AD hippocampal tissues and in cultured neuronal cells exposed to Aβ42.

### Blocking p53 activity in AD mice reverses formation of VDR/RXR complex and ameliorates AD pathology

Because both VDR and p53 are known to regulate autophagic activity (22-24) and the impaired autophagy flux is functionally linked to amyloid deposition and gliosis, we speculated whether or not the VDR/p53 complex might contribute to the impaired autophagic flux in AD. We performed an RNAi knockdown study of p53 and VDR in SH-SY5Y cells and found that knocking down either p53 or VDR could reverse the autophagy and apoptosis caused by Aβ42 treatment alone or in combination with vitamin D_3_ (Fig. 1*J* and SI Appendix, Fig. S6), suggesting that the VDR/p53 interaction played an important role in mediating the Aβ-induced autophagic neurodegeneration in AD. To validate this result *in vivo*, we treated AD mice with p53 inhibitor PFTα and found that it decreased their autophagic protein LC3II levels (Fig. 3*A* and 3*B*). We also measured S349-phosphorylated p62 (P-S349) levels because site-specific phosphorylation of p62 has been implicated in the disruption of autophagy-mediated protein degradation in AD brains(25). We found p53 inhibitor to have decreased P-S349 levels, which are normally increased in AD brain (Fig. 3*A*, second panel). Lysosomal dysfunction has been found to lead to accumulation of autophagosomes and neurodegeneration in AD (26). We also found that cathepsin B, a negative feedback regulator of lysosomal biogenesis (27), was concomitantly increased in brain tissues obtained from mouse brain (Fig. 3*A*, 4th panel). Together, these findings further indicate that VDR/p53 plays an important role in impairing autophagic flux in AD and suggest that blocking the pathway could potentially restore the impairment in autophagy. To further our understanding, we conducted a microarray analysis to profile the gene expression response to PFTα treatment in AD mice. Like our Western blot and immunohistochemistry assays, gene set enrichment analysis (GSEA) revealed that treatment with the inhibitor enriched autophagy-related genes in AD hippocampal tissues (Fig. 3*C*). A tight coupling between autophagy and inflammatory stress in AD has previously been reported (28, 29). Similarly, PFTα treatment effectively suppressed this inflammatory response (fig. S7). Considered together, these results suggest that the VDR-p53 pathway may contribute to autophagic neurodegeneration in AD.

**Figure 3.**
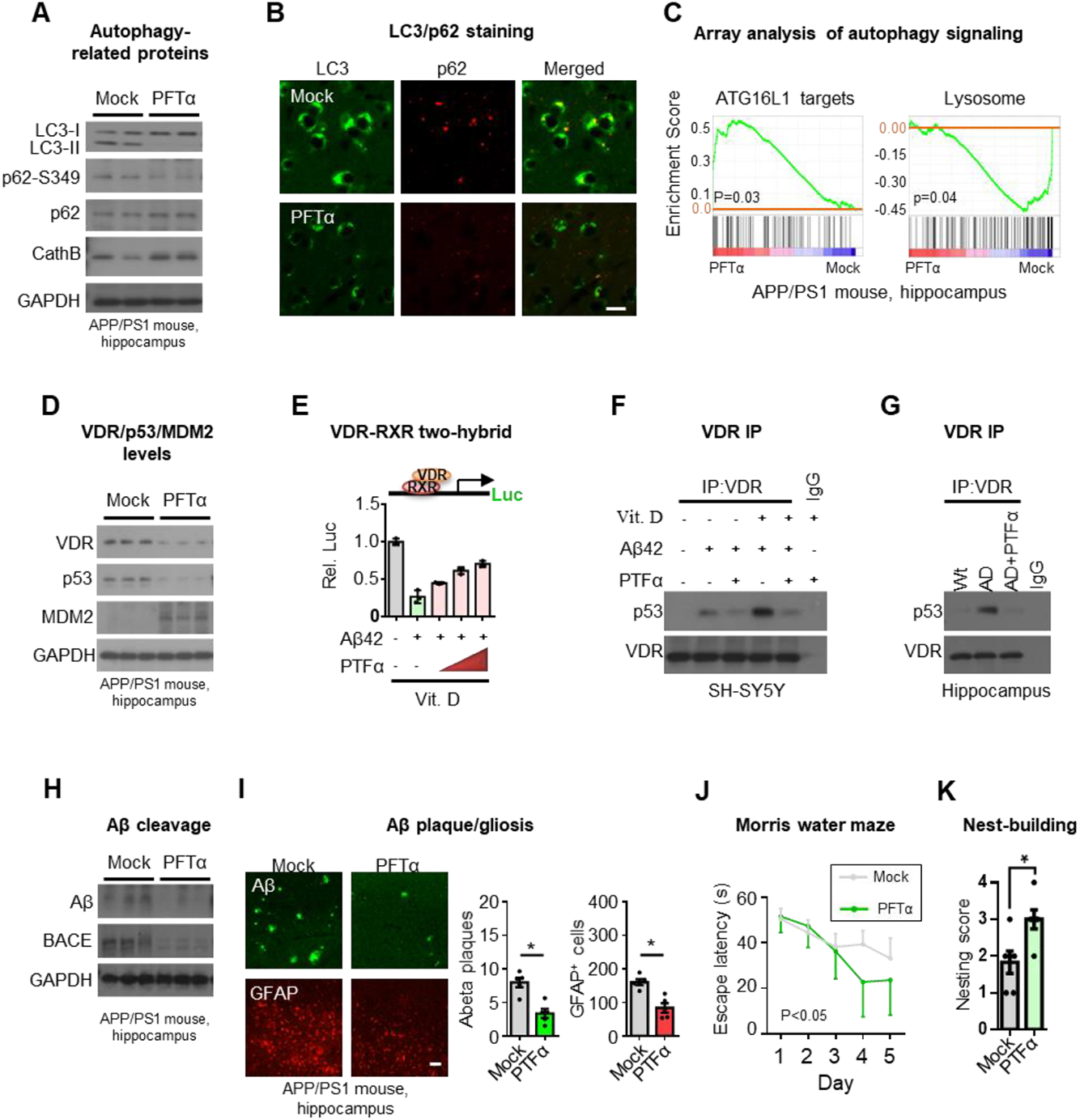
Reversal of brain pathology and cognitive function by inhibiting p53 activity in APP/PS1 AD mice. *(A)* Western blot analysis of autophagic markers LC3, p62, ser349 phosphorylated p62 (p62-S349), and lysosomal protease cathepsin B (cathB), in the hippocampal lysates of APP/PS1 mice. Four-month-old APP/PS1 mice raised on vitamin D_3_-sufficient diets were intraperitoneally injected weekly with 3 mg/kg of p53 inhibitor pifithrin-α (PFTα) for 8 months before harvesting hippocampal tissues for analysis. *(B)* Representative immunofluorescent micrographs of LC3 and p62 staining in the hippocampal sections of APP/PS1 mice. Scale bars, 5 μm. *(C)* Gene Set Enrichment Analysis (GSEA) showing significant functional gene sets differentially expressed in APP/PS1 mice after PFTα treatment, which include ATG16L1 (CADWELL_ATG16L1_TARGETS_UP) and lysosome (KEGG_LYSOSOME). The upregulation of ATG16L1 target genes refers to the reduction of ATG16L1 function. *(D)* Western blot analysis of VDR, p53 and MDM2 in the hippocampal lysates of APP/PS1 mice treated with or without p53 inhibitor. *(E)* Mammalian two-hybrid assays for studies of interaction of VDR with RXR in neuronal cells exposed to PTFα. *(F)*-*(G)*, Western blot analysis of co-immunoprecipitation of VDR/p53 complex in SH-SY5Y cells (F) and in the hippocampus lysate of APP/PS1 mice (G) with the indicated treatments. *(H)* Western blot analysis of Aβ and BACE levels in the hippocampal lysates of APP/PS1 mice treated with or without p53 inhibitor. *(I)* Representative immunofluorescent micrographs of amyloid aggregates (anti-Aβ) and gliosis (anti-GFAP) double staining in the hippocampal tissue sections of APP/PS1 mice treated with or without p53 inhibitor (left panel). The average percentage of surface area with Aβ plaques and gliosis in five consecutive sections of hippocampus per animal (n=5) was quantified by ImageJ and presented as the mean ± SEM (right panel). Significantly different from control group at *P < 0.05 by unpaired-sample t-test. Scale bars, 20 μm. *(J)*-*(K)* Cognitive performance assays for the AD mice treated with p53 inhibitor. APP/PS1 mice were given with or without weekly injections of PTFα (n=6 mice) starting at the age of 4-month. APP/PS1 mice at 12-month of age were used for the Morris Water Maze test (J) or Nest construction behavior (K). *P<0.05 by ANOVA for the Morris Water Maze assays. Unpaired-sample t-test was used to compare activity of nest construction behavior.

Finally, since the impaired autophagic signaling was effectively rescued by the p53 inhibitor, we performed an experiment to determine whether the AD brain’s molecular and morphological changes as well as the cognitive decline could also be ameliorated with the use of p53 inhibitor. We found that it not only decreased protein levels of p53 and VDR (Fig. 3*D*) but it also concomitantly increased the levels of MDM2 (Fig. 3*D*, 3ed panel), both with or without vitamin D_3_ supplementation. It should also be noted that the disrupted VDR/RXR interaction was also found to be restored (Fig. 3*E*) following p53 inhibitor treatment (Fig. 3*F* and 3*G*), suggesting that disrupting the VDR/p53 complex could reverse the binding of VDR to RXR. Based on these findings, we believed we would also be able to observe improvement in brain lesions. Indeed, Western blot and immunohistochemistry studies both revealed significant attenuation in Aβ deposits, BACE activity, reactive gliosis, and neuronal apoptosis in the AD brains, regardless of vitamin D_3_ supplementation (Fig. 3*H*, 3*I* and SI Appendix, Fig. S8). Finally, we wanted to know whether cognitive functioning and performance behavior would also be improved by treatment with p53 inhibitor. Mice that were given the p53 inhibitor showed significant improvement in both the Morris Water Maze test and nest construction behavior (Fig.3*j* and 3*k*). These results suggest that VDR-p53 pathway might be targeted therapeutically in the treatment of AD.

### APP/PS1 AD mice given vitamin D sufficient diet exhibit decreased serum vitamin D levels

Because the genomic action of VDR was found to be readily impaired in AD brain as noted above, it was imperative to clarify whether there is a causal link or whether vitamin D deficiency is just a disease consequence of AD. To test this, we fed both APP/PS1 and wild-type (WT) mice a vitamin D_3_ sufficient diet (600 IU/Kg of cholecalciferol) and measured their serum vitamin D levels during the early stages of life. Intriguingly, beginning as early as four months, the AD mice started exhibiting significantly lower serum 25(OH)D_3_ levels compared with those two-month-old. There was no difference in serum vitamin D levels in WT over the study period (Fig. 4*A*). These results may suggest that vitamin D deficiency to be the consequence of AD, rather than being caused by the lack of vitamin D in the diet.

**Figure 4.**
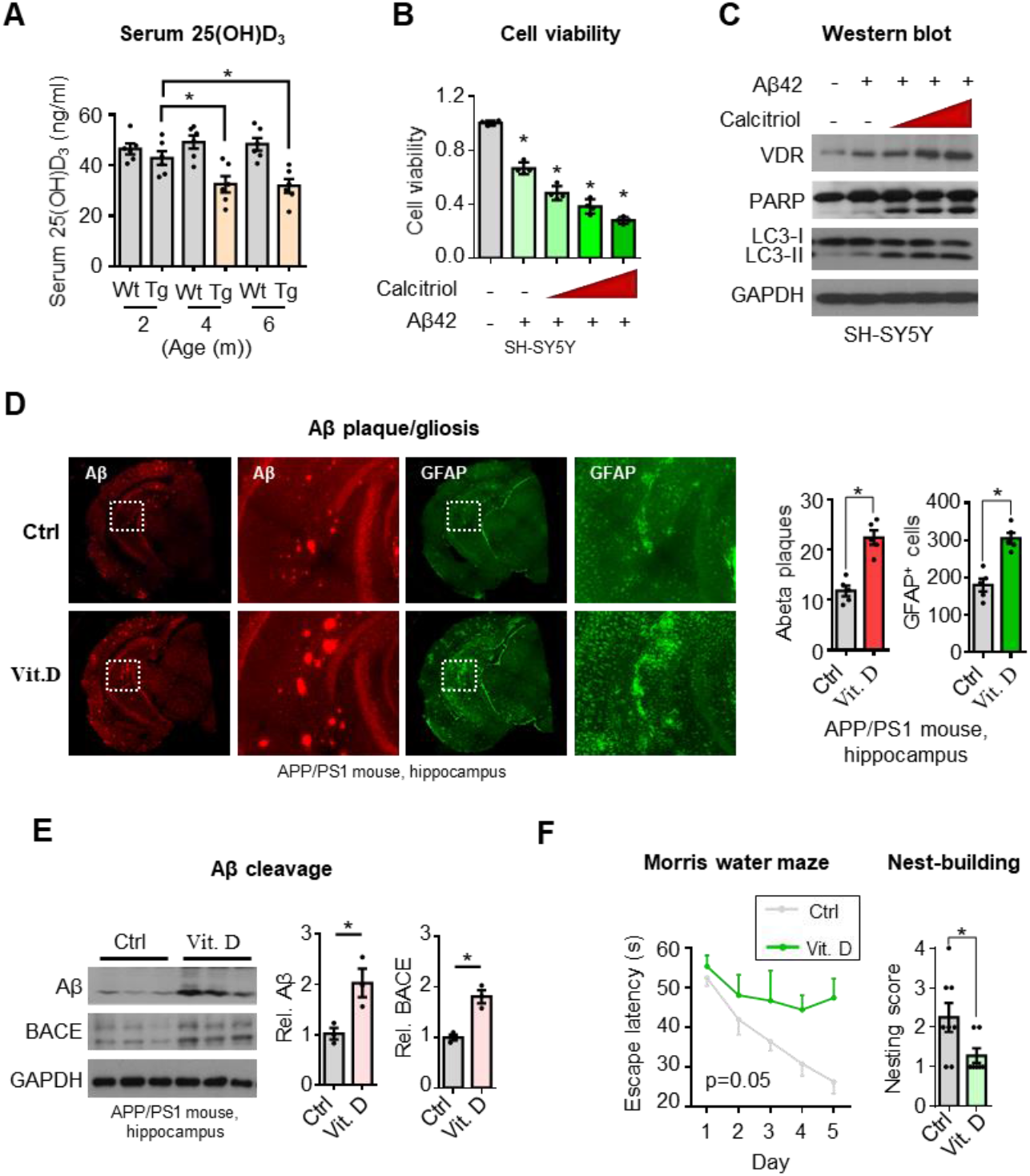
Aggravated AD pathology in APP/PS1 AD mice supplemented with dietary vitamin D_3_. *(A)* Serum 25(OH)D_3_ levels in APP/PS1 and wild-type mice. Mice were weaned at 4-weeks of age (±3 days) and maintained on a vitamin D_3_ sufficient diet (600 IU/Kg of cholecalciferol). Serum vitamin D3 levels were determined by 25-(OH) vitamin D3 enzyme-linked immunosorbent assay (EMSA) at the indicated time points (n=5). Results are shown as mean ± SEM. *P<0.05 by unpaired t-test. *(B)* Vitamin D_3_ supplementation causes a dose-dependent decrease of cell viability in SH-SY5Y cells exposed to β-amyloid. SH-SY5Y cells were treated with or without 4 μM of Aβ42 plus 0 μM, 1 μM, 3 μM or 10 μM cholecalciferol for 6 h prior to assays. WST cell viability assays were used to determine apoptotic cell death. Values are represented as the mean ± SEM and *P< 0.05 by unpaired t-test. *(C)* Western blot analysis of VDR, apoptotic and autophagic marker proteins in neuronal cells exposed to Aβ42 or plus without or without Vitamin D_3_. SH-SY5Y cells were treated with 4 μM Aβ42 alone or or Aβ42 plus 10, 30, or 100 nM calcitriol for 6 h before harvesting cell lysates for analysis. *(D)* Representative immunofluorescent micrographs of gliosis (anti-GFAP, GA5) and amyloid aggregates (anti-Aβ, D54D2) in cortex and hippocampal tissues of APP/PS1 mice. Four-month-old APP/PS1 mice were fed with vitamin D_3_-supplemented (0.8 mcg cholecalciferol/day, Vit. D) or vitamin D_3_-sufficient diets (0.06 mcg cholecalciferol/day, Ctrl) for 2 months before harvesting brain tissues for analysis. Sections of cortex or hippocampus were stained with the indicated antibodies. The average percentage of surface area with Aβ plaques in five consecutive sections per animal (n=4-7) was quantified by ImageJ in right panel. Significantly different from control group at P < 0.05 by unpaired t-test. *(E)* Western blot analysis of Aβ production and β-secretase 1 (BACE1) levels in hippocampal lysates of APP/PS1 mice supplemented with or without vitamin D_3._ Densitometrical quantification of Aβ and BACE bands were normalized to GAPDH. *P < 0.05 by unpaired t-test (right panel). *(F)* Cognitive performance for AD mice supplemented with vitamin D_3_. Four-month-old APP/PS1 mice were fed with vitamin D_3_–fortified (0.8 mcg cholecalciferol/day, Vit. D) or vitamin D_3_-sufficient diets (0.06 mcg cholecalciferol/day, Ctrl) for 8 months before Morris Water Maze (left panel) or nest construction behavior testing (right panel). Data of Morris Water Maze were analyzed by ANOVA (*P<0.05) and nest construction behavior by unpaired t-test.

### Faster disease progression after vitamin D supplementation in AD mice

Finding that the VDR-RXR heterodimer for transducing the genomic vitamin D signal is impaired in AD led us to question the seemingly common assumption that vitamin D supplementation may exert protective effect of on AD (30). We first assessed the potential impact of vitamin D supplementation on Aβ42-treated neuronal cell line. The results showed that the incubation of vitamin D (calcitriol or calcidiol) with SH-SY5Y cells being exposed to Aβ42 resulted in a significant dose-dependent increase in apoptosis and autophagy (Fig. 4*B*, 4*C* and SI Appendix, Fig. S9), suggesting a potential damaging effect of vitamin D on neuronal cells being exposed to Aβ42. Encouraged by this result, we proceeded to determine whether vitamin D supplementation could exert a similar detrimental effect on AD progression. Adding 0.8 mcg/day of cholecalciferol (a pro-hormone form of vitamin D_3_) to the diets of these APP/PS1 AD mice for two months resulted in more severe Aβ plaque deposits, reactive gliosis, and neuronal apoptosis in the hippocampus and cortex regions compared to controls (Fig 4*D*). Western blot analyses also revealed increased levels of pro-degenerative factors, including Aβ, β-secretase 1 (BACE1), Nicastrin (a subunit of the γ-secretase complex), TNFα, and autophagy in the hippocampal lysates of experimental mice (Fig. 4*E* and SI Appendix, Fig. S10).

Both groups of mice were also subjected to the Morris water maze test and nest construction test. Those administered vitamin D displayed worse cognitive functioning and performance behavior than the controls (Fig. 4*F*). These results of these *in vivo* and *in vitro* experiments suggest that over-supplementation of vitamin D can exacerbate AD neurodegeneration.

## Discussion

This study found evidence of a reverse causal link between vitamin D deficiency and AD, indicating that vitamin D deficiency may be an early disease manifestation in AD and not a cause. While the vitamin D levels were found to be low in both human and mouse with AD, it is intriguing to find that VDR were inversely activated in the brains of AD mice and patients. Other studies report findings that may also support this notion. Older African-Americans are two to three times more likely to develop AD than elderly whites (31, 32), while the African-Americans had higher mean VDR levels (32-34) but much lower serum vitamin D concentrations (35). Another example of the converse relationship between vitamin D concentration and VDR levels has been reported in patients with insulin resistance and obesity, who have been found have deficient levels of vitamin D on the one hand but increased levels of VDR in adipose tissue on the other (36). Notably, in addition to AD, patients with vascular disease, thyroid disorders, and osteoporosis are most likely to have decreased levels of serum vitamin D and, of course, be at higher risk for dementia (37, 38). It is therefore logical that future studies should explore whether the decrease of vitamin D is actually a common pathological response that occurs in many aging-associated diseases.

The inverse correlation of VDR protein levels and the concentration of its canonical ligand vitamin D in serum suggests that VDR could be activated by a non-genomic mechanism. Thus, we explored how the VDR proteins can be upregulated in the AD brains. In our exploration, we found the genomic VDR-RXR signaling pathway to be compromised by Aβ in AD, leaving the VDR/p53 complex to form in the cytosol and cause damage to AD brains. To determine whether the non-genomic activation of VDR/p53 played a role in AD, we used a chemical inhibitor of p53 to block the negative activity of VDR/p53 and reverse the brain pathology and cognition impairment in the AD mouse model. This suggests that the VDR/p53 signaling could potentially be used as a therapeutic target in the treatment of AD. However, use of p53 inhibitor could potentially increase the risk of tumor development because it is a key tumor suppressor protein. Consequently, further studies are needed to identify upstream modulators or downstream effectors for VDR-p53 signaling to avoid potential pitfalls.

Vitamin D deficiency in early childhood is linked to impaired neurodevelopment and skeletal health, and the most effective and accepted approach to resolve this issue is through vitamin D supplementation (39). The same deficiency in seniors is also considered as a common health risk factor that affects dementia, cardiovascular diseases, diabetes, cancers and several other chronic illness and geriatric syndromes. However, the need for its supplementation to prevent these diseases is currently debatable (40, 41). If deficiency of vitamin D is not a causal factor for AD and the genomic pathway of vitamin D has been compromised, these would lead one to doubt whether vitamin D repletion strategy could protect against AD. Indeed, the findings of our animal experiments suggested that the prolonged vitamin D supplementation actually exacerbates AD. Supporting this, our preliminary study of a nationwide longitudinal study revealed over-supplementation with vitamin D could promote the progression of dementia (unpublished data). These results suggest that caution be exercised in taking vitamin D3 supplements by AD patients. In line with this, multiple recent randomized clinical trials have also demonstrated no genuine health benefits of correcting vitamin D deficiency to reduce cardiovascular disease, type 2 Diabetes and chronic kidney disease (40). It is therefore clear that, until large careful clinical trials prove otherwise, older adults with dementia should be discouraged from taking vitamin D supplements. The results of our mechanistic analysis are important as we try to clarify why supplementation with vitamin D may not be the best way to address vitamin D deficiency with AD. Our findings, however, do not exclude a potential benefit of vitamin D supplementation for lowering AD risk in normal subjects when the neuroprotective VDR-RXR pathway remains intact.

## Materials and Methods

### Human brain tissue and ethics statement

All human brain tissues were obtained from the Brain and Tissue Bank at the University of Maryland. In total, 58 brain tissues were used. Among them, 40 were from individuals with a clinical diagnosis of probable AD, which includes 21 men and 19 women, with an average age of 80.1 ± 8.8 years, and postmortem interval of 10.25 ± 6.7 hours. Another 18 brain tissues were from individuals without neurological disorders, which include 11 men and 7 women, with an average age of 74.1±6.9 years, and a postmortem interval of 14.9 ± 8.6 hours. All studies and protocols were approved by the Research Ethics Committee at National Health Research Institutes (approved protocol no. EC1001103).

### Mice

Double transgenic APP/PS1 mice (Cat# 037565-JAX, RRID:MMRRC_037565-JAX) were purchased from Jackson Laboratory (Bar Harbor, ME, USA) to breed with wildtype B6C3F1/Bltw (C57BL/6N background) mice. Mice were weaned at 4-weeks of age (±3 days) and fed with the subnormal dosage of vitamin D_3_ diet (600 IU/Kg of cholecalciferol, corresponding to an intake of 0.06 mcg/day) (42, 43). Since it has been reported that feeding mice with 1-2 mcg of cholecalciferol per day for 12 weeks can significantly increase serum 25(OH)D_3_ but do not influence serum calcium level (44), APP/PS1 mice at 4 months of age were divided randomly into experimental group and control group. The mice of the experimental group were fed with 0.8 mcg of cholecalciferol per day (as D_3_-supplemented diet; Research Diets, Inc; Match Altromin 1320 With 8044 IU Vitamin D3/kg, Cat# D13031002) and control groups with 0.06 mcg per day (control diet; Altromin 1320 diet; Cat# Altromin 1320) for 2 to 12 months before assays. To block p53 activation in AD, the four-month-old APP/PS1 mice under vitamin D_3_-sufficient diet condition were intraperitoneally injected weekly with 3 mg/kg of p53 inhibitor Pifithrin-α (PFTα, Sigma-Aldrich, Cat# P4359) for 3 months before harvesting hippocampal tissues for analysis. For the Morris water maze test and nest construction behavior assays, PFTα was given to AD mice for 8 months. Serum 25(OH)D_3_ levels in APP/PS1 and wild-type mice were determined by Vitamin D_3_ EIA Kit (Cayman Chemical, Cat# 501050) at the indicated time points. All experimental animal procedures and protocols were approved by the Institutional Animal Care and Use Committee at NHRI (approved protocol no. NHRI-IACUC-101057-A and NHRI-IACUC-103136-A).

### Antibody

The antibodies used in this study are listed as follows: VDR (C20), Santa Cruz Biotechnology, Cat# sc-1008, RRID:AB_632070; VDR(D6), Santa Cruz, Biotechnology

Cat# sc-13133, RRID:AB_628040; GAPDH, GeneTex, Cat# GTX100118, RRID:AB_1080976; Tubulin, GeneTex, Cat# GTX112141, RRID:AB_10722892; PARP-1/2 (H-250), Santa Cruz Biotechnology, Cat# sc-7150, RRID:AB_216073; LC3B, Cell Signaling Technology, Cat# 4108, RRID:AB_2137703; Beta-Amyloid-1-16 antibody, BioLegend, Cat# 803014, RRID:AB_2728527; β-Amyloid Antibody, Cell Signaling Technology, Cat# 2454, RRID:AB_2056585; BACE (M-83), Santa Cruz Biotechnology, Cat# sc-10748, RRID:AB_2061505; GFAP (Clone SP78), MybioSourse, Cat# MBS302899, DISCONTINUED; GFAP (GA5), Cell Signaling Technology, Cat# 3670, RRID:AB_561049; p53 (DO-1) Santa Cruz Biotechnology Cat# sc-126, RRID:AB_628082; Cathepsin B Antibody (FL-339), Santa Cruz Biotechnology, Cat# sc-13985, RRID:AB_2261223; Phospho-SQSTM1/p62 (Ser349), Cell Signaling Technology, Cat# 95697, RRID:AB_2800251; SQATM1/p62 (GT1478),

Thermo Fisher Scientific, Cat# MA5-27800, RRID:AB_2735371; TNF-α (D2D4) XP® Rabbit mAb, Cell Signaling Technology, Cat# 11948, RRID:AB_2687962; Lamin B (M-20) antibody, Santa Cruz Biotechnology, Cat# sc-6217, RRID:AB_648158; Alexa 488 chicken anti-rabbit IgG(H+L), Thermo Fisher Scientific, Cat# A-21441, RRID:AB_2535859; Alexa 594 chicken anti-mouse IgG(H+L), Thermo Fisher Scientific, at# A-21201, RRID:AB_2535787; Alexa 594 chicken anti-rabbit IgG(H+L), Thermo Fisher Scientific, Cat# A-21442, RRID:AB_2535860; Alexa 488 chicken anti-goat IgG(H+L), Thermo Fisher Scientific Cat# A-21468, RRID:AB_2535871; Peroxidase-AffiniPure Goat Anti-Rabbit IgG (H+L), Jackson ImmunoResearch Labs, Cat# 111-035-144, RRID:AB_2307391; Peroxidase-AffiniPure Goat Anti-Mouse IgG (H+L), Jackson ImmunoResearch Labs, Cat# 115-035-146, RRID:AB_2307392; Peroxidase-AffiniPure Rabbit Anti-Goat IgG (H+L), Jackson ImmunoResearch Labs, Cat# 305-035-003, RRID:AB_2339400; Mouse anti-Rabbit light chain; HRP conjugate, Millipore, at# MAB201P, RRID:AB_827270; HRP-conjugated AffiniPure Mouse Anti-Rabbit IgG Light Chain, Bclonal Cat# AS061, RRID:AB_2864055; HRP-conjugated AffiniPure Goat Anti-Mouse IgG Light Chain antibody, ABclonal, Cat# AS062, RRID:AB_2864056; Duolink In Situ PLA Probe Anti-Rabbit PLUS antibody, Sigma-Aldrich, Cat# DUO92002, RRID:AB_2810940; Duolink In Situ PLA Probe Anti-Mouse MINUS Antibody, Sigma-Aldrich, Cat# DUO92004, RRID:AB_2713942.

### Cell Culture

The following cell lines were used in this study: SH-SY5Y (human neuroblastoma, ATCC CRL-2266), IMR-32 (human neuroblastoma, ATCC CCL-127) and MPNC (mouse primary neural lineage cells, a gift from Dr. Hsing I Huang at Chang Gung University). SH-SY5Y and IMR32 were cultured in MEM (Invitrogen, USA) and MPNC in DMEM (Invitrogen, USA). Cells were grown at 37°C in a 5% CO2 humid atmosphere.

### Cells transfection, mammalian two-hybrid assay and Cyp24a1 reporter assay

Oligomeric β-amyloid (Aβ42, MDbio, Inc) was prepared as described previously (45). RNAi-mediated knockdown of VDR or p53 was performed by transient transfection of siVDR, sip53, or a control siRNA (Stealth siRNA, Invitrogen, USA) into SH-SY5Y cells with DharmaFECT (Dharmacon, USA) at a concentration of 50 nM in 6-well culture plates. Transient overexpression of VDR in SH-SY5Y cells was performed by transfecting a VDR expression plasmid (pcDNA3-VDR) or a mock control plasmid (pcDNA3) with Lipofectamine 2000 (Invitrogen, USA). For mammalian two-hybrid assays to assess the interaction between VDR and RXR, SH-SY5Y cells were co-transfected with 100 ng of pCMV-BD-RXRα (as bait), 100 ng of pCMV-AD-VDR (as prey), 500 ng of pFR-luc (as a reporter) and a control plasmid (pRL-null constitutively expressing low levels of Renilla reniformis) in 6-well plates as described in Dr. Jurutka’s publication (46) with minor modifications. The transfected cells were then treated with Aβ42 (1-4 μM) or calcitriol (100 nM) for 6 h before luciferase activity assay (Dual-Luciferase Reporter Assay, Promega). A 587-bp region of the human Cyp24a1 gene promoter that contains two VDRE motifs was cloned into the promoter-less luciferase expression vector pGL3-basic (Promega) (47). The transfected cells were then treated with Aβ42 (1-4 μM) or calcitriol (100 nM) for 6 h before Dual-Luciferase Reporter Assay.

### In situ proximity ligation assay

A Duolink® PLA Starter Kits ((Sigma-Aldrich, St. Louis, MO) was used to detect in situ proximity ligation assay for VDR/p53 interactions in postmortem brain tissues. Paraffin-embedded human brain sections were incubated with mouse anti-p53 (DO-1) and rabbit anti-VDR (C20) antibodies (Santa Cruz Biotech) in antibody diluent buffer overnight at 4°C, followed by incubation with Duolink anti-mouse MINUS and anti-Rabbit PLUS secondary antibodies for 1h. For detection, the Duolink in situ detection reagent-RED was used.

### Cell viability and TUNEL staining

Colorimetric WST-1 assay (Roche) was used to determine cell viability. The absorbance was measured by a spectrophotometer (SpectraMax Plus from Molecular Devices) at 450 nm against a reference at 690 nm. The optical density values relative to the control cells in the assay represent the percentage of viable cells. To detect cell apoptosis, the TUNEL assay was performed using the ApoAlert™ DNA Fragmentation Assay Kit (Clontech) to detect the presence of DNA fragmentation in frozen tissue sections. The fixed sections were washed twice with PBS before incubating in the permeabilization solution (0.2% Triton X-100 in PBS) on ice for 10 min. The sections were washed twice in PBS and then incubated in TUNEL reaction mixture at 37°C in the dark in a humidified atmosphere for 1 h. The stained sections were washed once again with PBS before mounting with DAPI mounting medium (VECTASHIELD) for fluorescence microscopy analysis.

### Quantitative real-time reverse transcription-PCR

Total RNA from brain tissues or culture cells were extracted using the illustra RNAspin Mini RNA Isolation Kit (GE Healthcare Life Sciences) for reverse transcription with the High-Capacity cDNA Reverse Transcription Kits (ABI Applied Biosystems, USA) according to the manufacturer’s instructions. The quantitative real-time Reverse Transcription-PCR (qPCR) analysis was performed using the Fast SYBR Green Master Mix (ABI Applied Biosystems, USA). Results were determined using respective standard curves calculations. The primers used in this study are listed as follows: Human VDR-F: 5’-CGA CCC CAC CTA CTC CGA CTT-3’; Human VDR-R 5’-GGC TCC CTC CAC CAT CAT TC-3’; Mouse VDR-F: 5’-GGA GCT ATT CTC CAA GGC CC -3’; Mouse VDR-R: 5’-GGG TCA TCG GAG CCT TCT TC-3’; Human GAPDH-F: 5’-CCT GCC AAA TAT GAT GAC ATC AAG-3’; Human GAPDH-R: 5’-ACC CTG TTG CTG TAG CCA AA-3; Mouse GAPDH-F: 5’-AAG GTC ATC CCA GAG CTG AA-3’; Mouse GAPDH-R: 5’-CTG CTT CAC CAC CTT CTT GA-3’; Human Cyp24a1-F: 5’-CAT CAT GGC CAT CAA AAC AAT-3’; Human Cyp24a1: 5’-GCA GCT CGA CTG GAG TGA C-3’.

### Microarray and GSEA gene sets analysis

The microarray of mouse Clariom S Assays (Thermo Fisher Scientific) for whole-transcript expression analysis were used in this study. We compared the gene expression profiles of hippocampal tissues from the APP/PS1 mice treated with or without p53 inhibitor (Pifithrin-α, 3 mg/kg). To analyze the signaling pathways that were impacted by the p53 inhibitor, we performed Gene Set Enrichment Analysis (GSEA) (48, 49) for the C2-curated gene sets and H-hallmark gene sets with a size of >15 genes. P<0.05 was considered significant.

### Quantification and statistical analysis

Statistical information, including n (number of patients or mice), mean and statistical significance values, is indicated in the figure legends. None specific method was used to determine whether the data met assumptions of the statistical approach. Statistical significance was determined with Graphpad Prism 6 using the tests indicated in each figure. Data were considered statistically significant at p < 0.05.

## Acknowledgments

We think UMB Brain and Tissue Bank (University of Maryland, Baltimore) for providing the human brain tissues used in this study. We thank Dr. Peter Jurutka for providing the pCMV-BD-RXRa, pCMV-AD-VDR, pRL-Null and pFR-luc plasmids for mammalian two-hybrid assays used to determine VDR-RXR interaction. We thank NHRI Optical Biology Core, Microarray Core Laboratory, and Animal Behavior Core Facility for technical supports. We also thank James Steed and Duane S. Juang for English editing of the manuscript. Funding: This work was supported by MOST 105-2321-B-400-00 2-MY3 and NHRI MG-104-PP-02.

