## Supplementary Information for "The non-genomic vitamin D pathway links β-amyloid to autophagic apoptosis in Alzheimer’s disease" (from another version of this preprint: version 1, 2021-05-07)

Jyh-Lyh Juang

**This PDF file includes:**

Figures S1 to S10

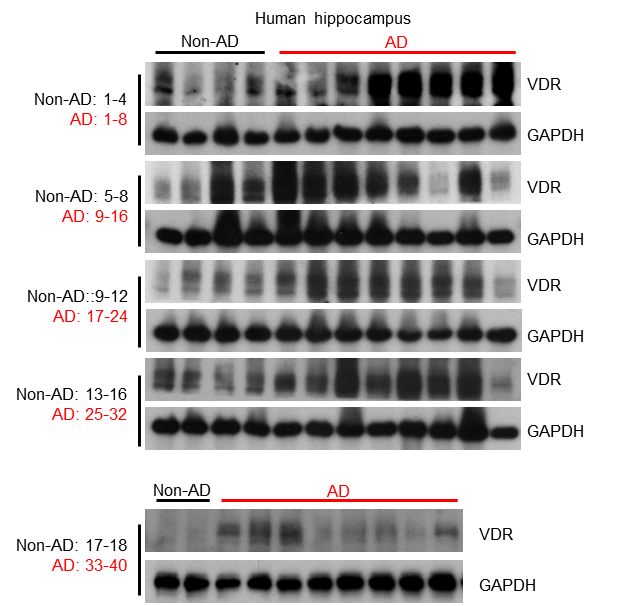

Fig. S1. Elevated VDR protein levels in hippocampus tissues of AD patients. Western blot analysis of VDR levels in hippocampus tissues of AD (n=40) patients and age and gender-matched non-AD controls ( n=18).

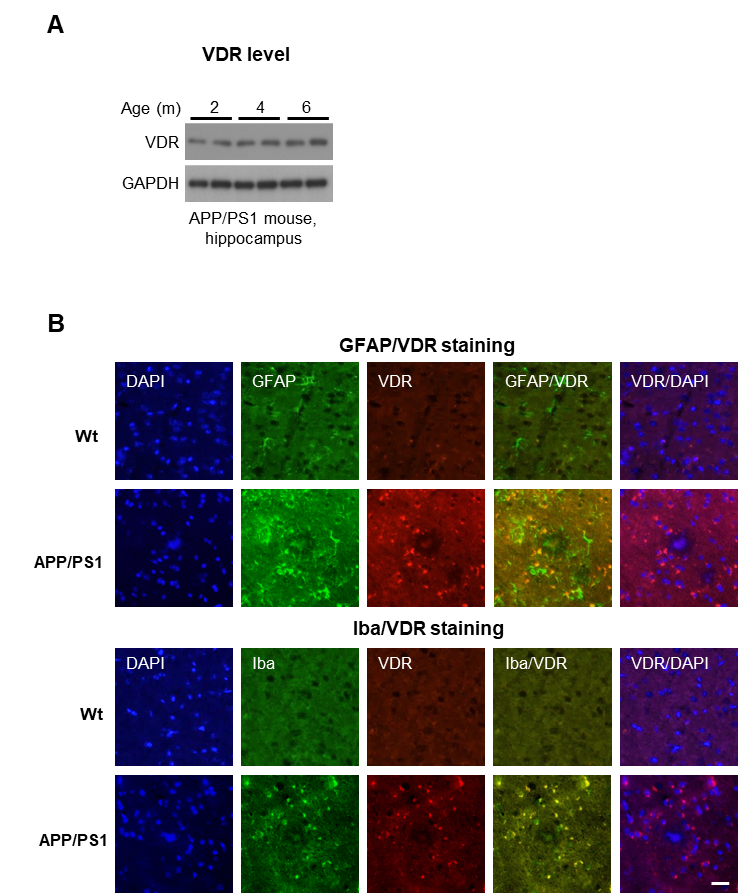

Fig. S2. Elevated VDR protein levels in hippocampus tissues of APP/PS1 mice. *(A)* Western blot analysis of VDR levels in the hippocampal lysates of APP/PS1 mice. *(B)* Representative images of immunohistochemistry analysis for GFAP (astrocytes marker), Iba (microglia marker), and VDR in sections of Aβ plague region of the hippocampus in APP/PS1 mice. Scale bars, 20 μm.

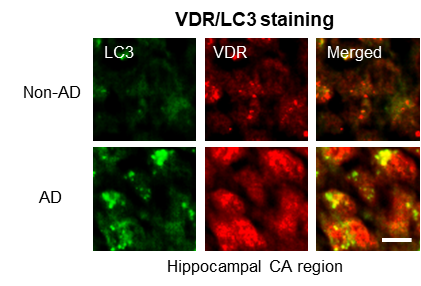
Fig. S3. Representative immunofluorescent micrographs stained for VDR and LC3 in the hippocampal tissue sections of human AD brains. (A) Aβ plague regions in human and mouse AD brains. Scale bars, 10 μm.

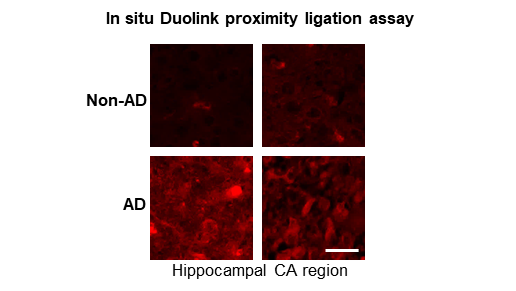
Fig. S4. Detection of VDR-p53 interaction in hippocampal tissues of AD patients by in situ Duolink proximity ligation assay. Scale bars, 20 μm.

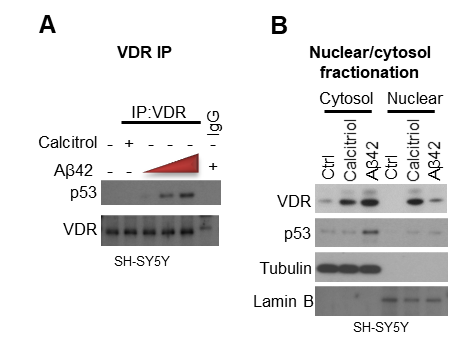
**Fig. S5. Switching the VDR’s heterodimerization partner from RXR to p53 by Aβ in SH-Y5Y neuron cells.** *(A)* Western blot analysis of co-immunoprecipitated VDR and p53 in neuronal cells. *(B)* Aβ42 increases cytosol p53. SH-SY5Y neuronal cells were treated with 100 nM calcitriol or 4 μM Aβ42 for 6 h prior to assay.

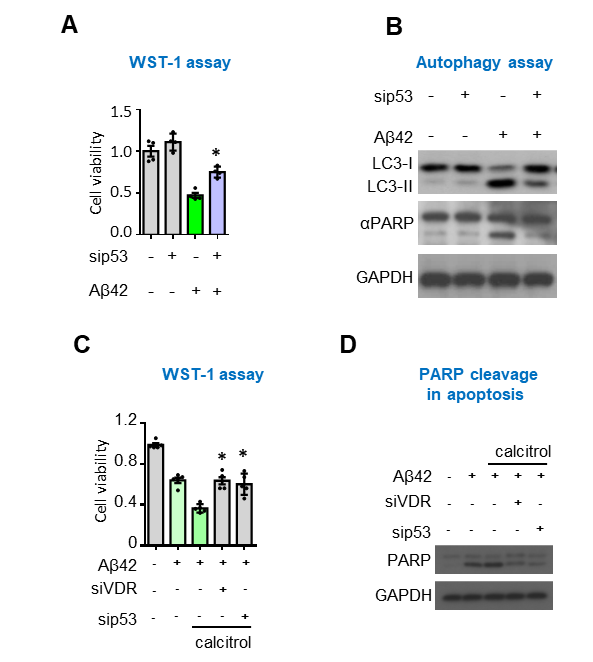
**Fig. S6. p53 modulation of the Aβ42-induced neuronal autophagy and apoptosis in neuronal cells.** *(A)* WST-1 assays for the Aβ42-induced apoptosis in SH-SY5Y cells after RNAi depletion of p53. *(B)* Western blot analysis of hallmarks of the Aβ42-induced autophagy and apoptosis in the p53-depleted SHS-Y5Y cells. *(C)*-*(D)* Additive enhancement of apoptosis by Aβ42 and calcitriol is ameliorated by RNAi depletion of p53. SH-SY5Y cells were treated with or without 4 μM of Aβ42 plus 100 nM calcitriol for 6 h prior to WST assays (C), or western blot analysis of PARP cleavage (D). Values are represented as the mean ± SEM and *P< 0.05 by unpaired t-test.

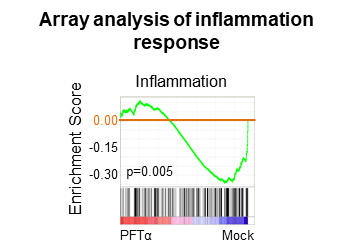
Fig. S7. Gene Set Enrichment Analysis (GSEA) showing inflammatory response gene sets differentially expressed in APP/PS1 mice after PFTα treatment. The inflammatory response base on the HALLMARK_INFLAMMATORY_RESPONSE gene set.

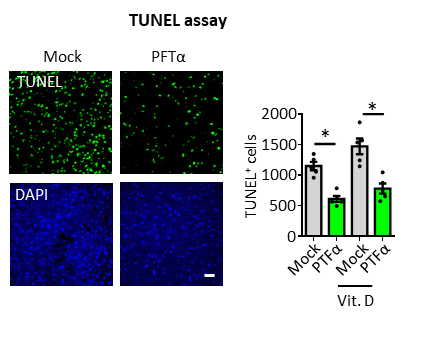
Fig. S8. p53 inhibitor amelioration of vitamin D3-aggravated Aβ aggregation and apoptosis. TUNEL immunofluorescent staining of brain sections in the APP/PS1 mice mentioned above. Scale bars, 50 μm. The TUNEL-positive signals in five consecutive sections per animal (n=5) were quantified by ImageJ and presented as the mean ± SEM (right panel). *P < 0.05 by unpaired t-test.

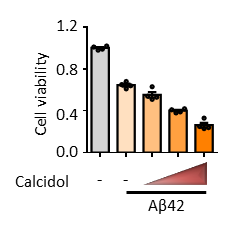
Fig. S9. Vitamin D_3_ supplementation causes a dose-dependent decrease of cell viability in SH-SY5Y cells exposed to β-amyloid. SH-SY5Y cells were treated with or without 4 μM of β-amyloid (Aβ42) plus 0 μM, 1 μM, 3 μM or 10 μM calcidiol (25-hydroxyvitamin D_3_) for 6 h prior to assays. WST cell viability assay was used to determine apoptotic cell death. Values are represented as the mean ± SEM.

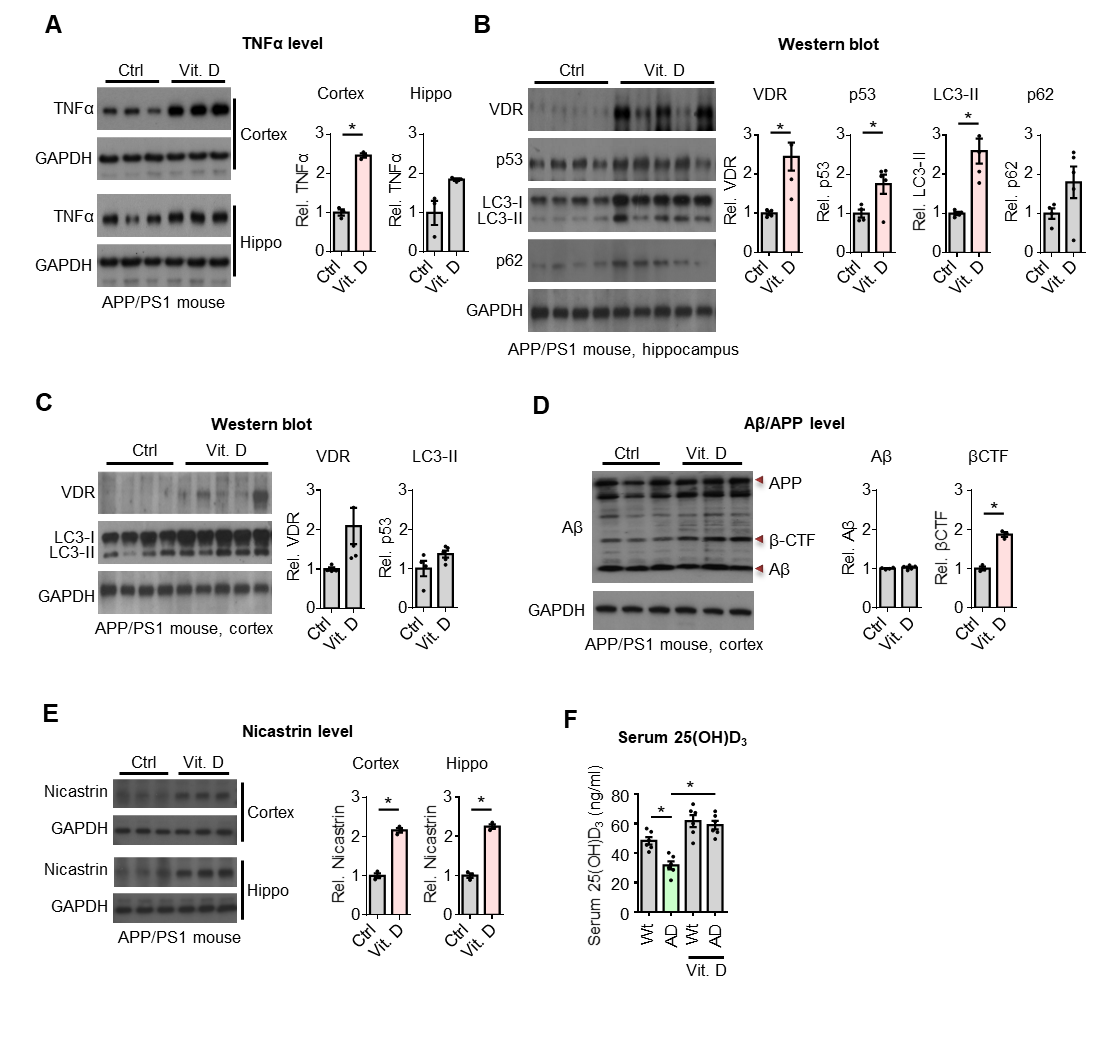
Fig. S10. Western blot analysis of inflammatory cytokine, autophagy- and APP processing-related proteins in the hippocampus and cortex of APP/PS1 mice supplemented with or without cholecalciferol. *(A)* The blots for lysates from the hippocampus and cortex were probed with antibodies against TNFα and GAPDH. Densitometrical quantification of indicated protein levels were calculated as ratios normalized with GAPDH. *P < 0.05 by unpaired t-test (right panel). *(B)* The blots for lysates from hippocampus were probed with antibodies against VDR, p53 and LC3. *(C)* The blots for lysates from cortex were probed with antibodies against VDR and LC3. *(D)* The blots for lysates from cortex were probed with antibodies against Αβ. *(E)* The blots for lysates from cortex were probed with antibodies against Nicastrin. *(F)* Cholecalciferol (25-hydroxyvitamin D3) supplementation restores serum 25(OH)D3 levels. Serum 25(OH)D3 levels were determined by Vitamin D3 EIA Kit and the results are shown as mean ± SEM. *P<0.05 by unpaired t-test.
